# Toxicity testing of two pesticides against the Carthusian snail *in vitro*

**DOI:** 10.1101/2023.09.01.555921

**Authors:** A.S. Bashandy, M.H Awwad

## Abstract

In this work, the land snail, *Monacha cartusiana*, was studied to assess the deadly toxic activity of two pesticides (lambda and methomyl) on this snail in *vitro*, utilizing contact and dipping techniques. The results revealed that methomyl was more effective against snails compared with lambda in both methods of treatment. However, the contact application strategy resulted in greater death rates than the poison dipping studies. Accordingly, the mortality rate was increased by increasing concentration, with contact technique which recorded 96.3% and 69.99±15.1% deaths by dipping way on concentration 3%. For this reason, the percent Lethal Concentration fifty (LC_50_) was 1.30% for one day and 1.67% for seven days for methomyl by contact and dipping techniques, respectively. On the other hand, the LC_50_ for lambda pesticide was achieved at 1.45% by contact method but was 7.51% by using the dipping application within ten days. As a result, mortality was greater after contact application than after dipping, and methomyl was disclosed to be more poisonous than lambda as a molluscicide.

## 1. Introduction

Terrestrial snails are one of the most serious threats to sustainable agriculture in many parts of the world (Barker, 2002). In Egypt, land snails are considered major pests of farms (Kassab and Daoud, 1964; El-Okda, 1979). Snail damage is usually caused by feeding and contaminating their bodies, feces, or slime, resulting in worse quality goods and diminished earnings. They may transmit illnesses or act as intermediary hosts for human and animal pathogens (Glesias *et al*., 2003; Heiba *et al*., 2018). *Monacha* species are considered a major agricultural pest, they cause a lot of harm to vegetable crops in various areas of Egypt (Wafaa *et al*., 2018).

Insecticides are used to control land snails that contaminate various crops. Thus, chemical pesticides are now the most efficient means of combating terrestrial Mollusca control. (El-Wakil and Radwan, 1991; Hanafy *et al*., 1998; Hussein *et al*., 1999; El-Khodary *et al*., 2001; Heiba *et al*., 2002; Genena, 2003; Abd El-All, 2004; Ismail *et al*., 2005 and Zedan *et al*. 2006). However, because some of these pesticides are ecologically highly stable, the danger of accumulation is quite significant, and as a result, their usage as insecticides has been limited (Ohayo *et al*., 1997). As a result, new and safe insecticides or molluscicides with distinct modes of action are required. The goal of this study is to evaluate the molluscicidal activity of two pesticides, Lambada and Methomyl, on the adult land snail, *Monacha cartusiana, in vitro*.

## 2. Materials and Methods

### 2.1. Pesticides that have been tested include

1. Lambada 5 % – Cyhalothrin E.C.
2. Methomyl (Lannate 90 % W.P) insecticide.

### 2.2. Land snail feeding and care

Adult *Monacha cartusiana* were obtained from polluted fields in the Kerdasa district of Giza Governorate, Egypt, during the spring of 2022. Snails were sent in sealed bags to the Malacology laboratory at the Dept. of Agric. Zoo. and Nema., Fac. of Agriculture, Al-Azhar University. Healthy individuals were placed in separate terrariums (30 x 20 x 15 cm) filled with moist soil and covered with muslin tied with a rubber band to prevent snails from escaping. Every day for two weeks, the animals were fed fresh lettuce leaves (*Lactuca sativa*). They were kept at a temperature of 20±2°C with the humidity level of the soil was 75% according to Mobarak 2003.

### 2.3. Valuation of sensitivity to adult Carthusian snails for two pesticides

Insecticides potential of methomyl (S-methyl-N-[(methylcarbamoyl) oxy] thioacetimide) compared with Lambda-Cyhalothrin as Molluscicidal were tested by preparing a series of concentrations were prepared using distilled water for two methods of contact and dipping techniques on tested species.

### 2.4. Toxicity test by contact technique

The contact toxicity of two pesticides was determined using the surface exposure technique, in which 2 ml of the various concentrations (0.5, 1, 1.5, 2, 2.5, and 3%) was dropped and spread over the bottom of a petri dish and then slowly rotating the dish in circles. The water disappeared after a few minutes at normal temperature, leaving just a tiny film of the tested chemicals at the applied concentration. Mourad (2014) subjected ten normal individual snails from the tested animals to the potential concentration of the tested compounds for 24 hour. Each of the treatments received three duplicates, in addition to an untreated control. Within 24 hour, dead snails were gathered up.

### 2.5. Toxicity test by dipping technique

Ten healthy snails were chosen for each replicate and deprived for 24 hour before the experiment began. The concentrations used have previously been mentioned by Lambada and Methomyl. The lettuce leaves were immersed in each concentration for one minute before being fed to the experimental animals. Then, for the next 10 days, untreated lettuce was introduced as needed for feeding. Untreated animals were fed lettuce that had simply been treated with water (Bashandy, 2018). Mortality percentages and half Lethal concentration of check pesticides were calculated after one day. and within ten days of treatment according to Finney (1952) using Bakr (2005) Computing program.

## 3. Results

Molluscicidal activity of Methomyl and Lambada against *M. cartusiana*. The collected data in Table (1) and Figure (1) displayed that, the high mortality ratio of methomyl and lambda was (96.33 and 90%) after a few hours of testing contact technique with a concentration of 3%, respectively. Also, the concentrations of 2 and 2.5% per 100 ml caused mortality percentages were (70 and 80%) and (60 and 70%) during one day. for methomyl and lambda, respectively. while the rest concentrations of pesticides recorded mortality under fifty percent of animals. Over and above, the Regression Linear Lethal concentration of the test animals for methomyl was higher than lambda with a rate of 1.3097%.

**Fig. 1:**
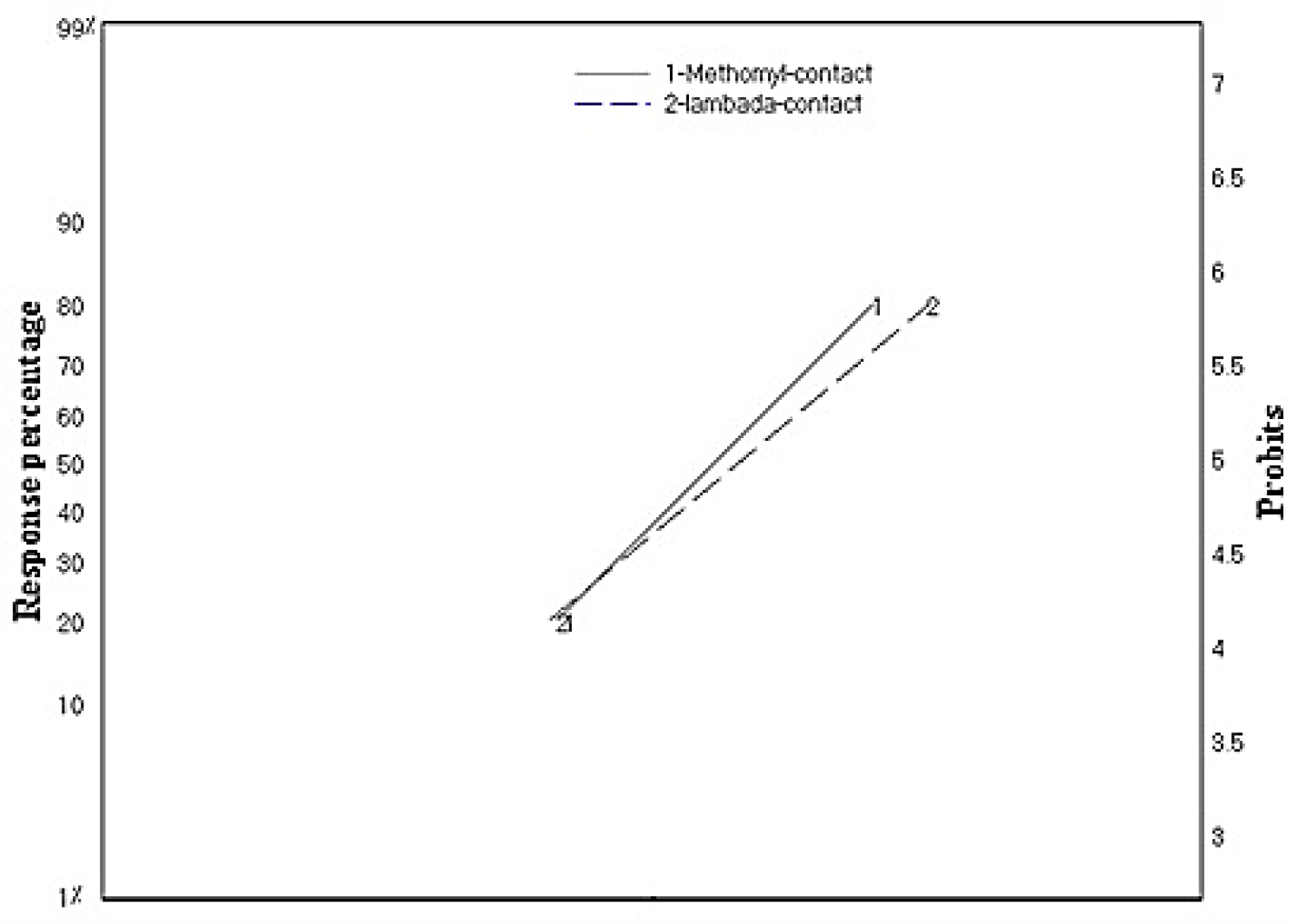
Regression Linear for pesticides by contact method contra *M. cartusiana* during 24 hrs.

**Table 1:** The contact technique was used to assess lambda and methomyl mortality in *M. cartusiana*.

| Pesticide | Con. (%/100 ml) | 0.5 | 1 | 1.5 | 2 | 2.5 | 3 | LC<sub>50</sub> (%/100 ml) |
| --- | --- | --- | --- | --- | --- | --- | --- | --- |
| Lambda 5% | Observed mortality % | 20 | 30 | 40 | 60 | 70 | 90.00 | 1.4513 (1.1779-1.7543) |
| Methomyl 90% |  | 20 | 30 | 40 | 70 | 80 | 96.33 | 1.3097 (1.0825-1.538) |

The lower and higher fiducial limits of 95% are provided in parentheses.

**Table 2:** The leaf dipping technique was used to assess lambda and methomyl mortality in *M. cartusiana*.

| Lambda 5% |  |  |  |  | Methomyl 90% |  |  |  |  |
| --- | --- | --- | --- | --- | --- | --- | --- | --- | --- |
| Con.<br>(%/ml) | Corrected mortality % per day |  |  |  | Con.<br>(%/ml) | Corrected mortality % per day |  |  |  |
|  | 1<sup>st</sup> | 3<sup>rd</sup> | 10<sup>th</sup> | mean |  | 1<sup>st</sup> | 3<sup>rd</sup> | 7<sup>th</sup> | mean |
| 0.5 | 3.33 | 6.67 | 10.00 | $6.66 \pm 1.5^e$ | 0.5 | 3.33 | 6.67 | 13.33 | $7.77 \pm 2.4^d$ |
| 1 | 6.67 | 10.00 | 13.33 | $10.00 \pm 1.5^{de}$ | 1 | 6.67 | 10.00 | 23.33 | $13.33 \pm 4.1^d$ |
| 1.5 | 10.00 | 13.33 | 16.67 | $13.33 \pm 1.5^{de}$ | 1.5 | 10.00 | 20.00 | 33.33 | $21.11 \pm 5.5^{cd}$ |
| 2 | 13.33 | 16.67 | 20.00 | $16.66 \pm 1.5^{cd}$ | 2 | 13.33 | 40.00 | 56.67 | $36.66 \pm 10.3^{cd}$ |
| 2.5 | 16.67 | 20.00 | 26.67 | $21.11 \pm 2.4^{bc}$ | 2.5 | 26.67 | 53.33 | 60.00 | $46.66 \pm 8.3^{bc}$ |
| 3 | 20.00 | 26.67 | 33.33 | $26.66 \pm 3.1^b$ | 3 | 33.33 | 83.33 | 93.33 | $69.99 \pm 15.1^{ab}$ |
| $LC_{50}$ | 13.87 | 13.19 | 9.43 | | $LC_{50}$ | 5.62 | 2.17 | 1.67 | |
| Control | 0.00<sup>a</sup> |  |  |  | Control | 0.00<sup>a</sup> |  |  |  |
| $LSD_{0.05}$ | 7.09 | | | | $LSD_{0.05}$ | 30.06 | | | |

Based on three replicates ± SEE, each plastic container has ten land snails.
Values in a column that are followed by the same letter are not statistically different at the 0.05 level.

The obtained data is in the table. (2) and figure. (2) disclosed that methomyl had high effects on land snails by leaf dipping method. It caused high mortality for individuals during seven days with moderate 69.99±15.1% by the last concentration (3%) with a lethal concentration fifty of snails was 1.67% per 100ml on regression linear. On the other hand, the pesticide lambda raised the percentage mortality by 26.66±3.1% for snails for ten days and LC_50_ was recorded at 9.43 % per 100ml on regression linear. But the lower concentration of pesticide lannate was higher than lambda which recorded mean mortality of 7.77±2.4 and 6.66±1.5% during interval periods respectively.

**Fig. 2:**
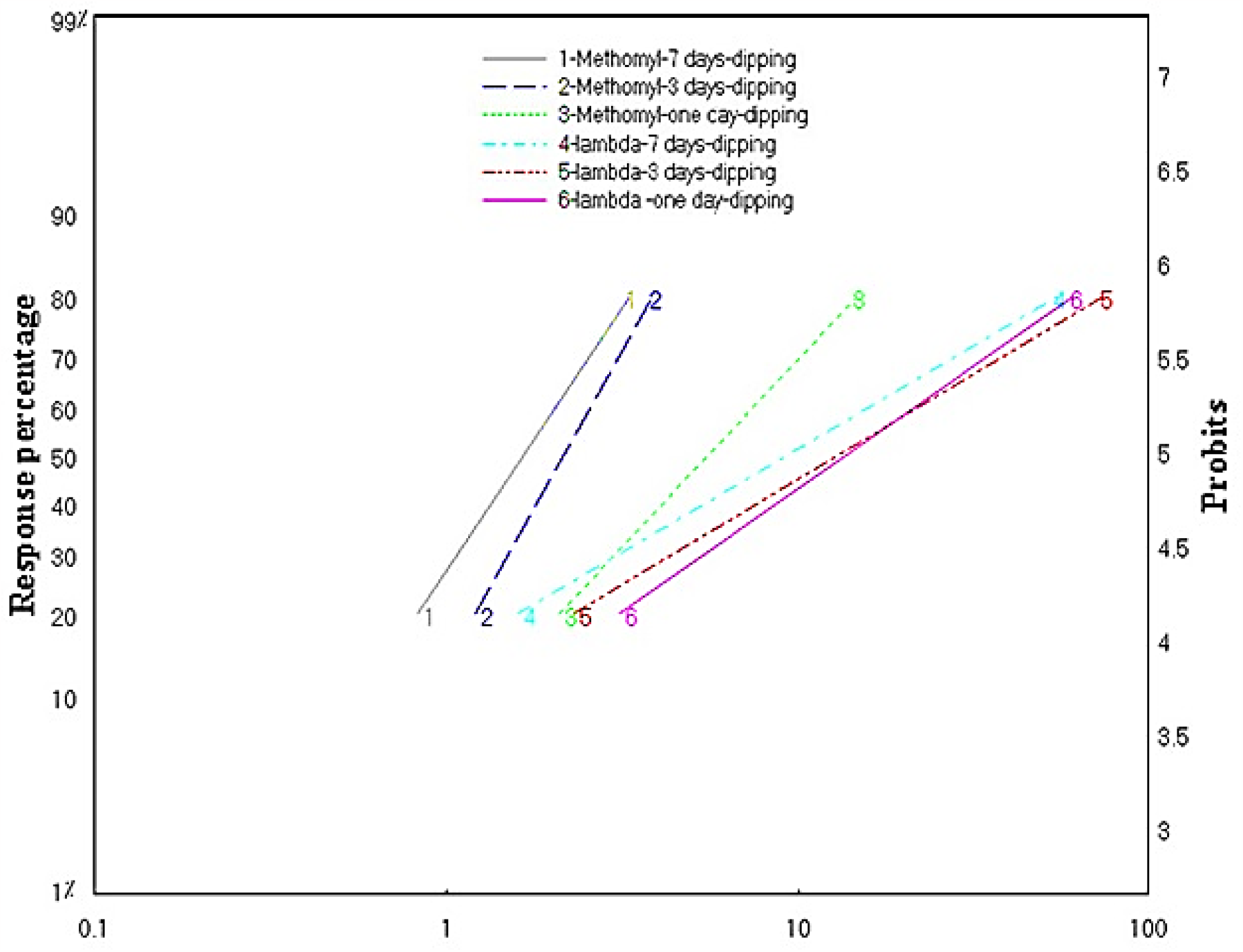
Regression Linear for pesticides by leaf dipping method contra *M. cartusiana* during exposure time.

The statical analyses for mean mortality percentage caused by each pesticide’s compounds proved, that there are significant between concentrations compared to the control that was treated with water only. There are significant differences between concentration 3% with the rest concentrations.

## 4. Discussion

Previous findings are consistent with those that came by Hammond, *et al*. (1996), who declare that the effectiveness of pesticides depends on their good application of them because the opposite leads to an increase in cost and impact on the environment. Pyrethroids, such as Lambda-cyhalothrin, damage the neurological system of insects, causing paralysis or death. (Vijverberg and van den Bercken, 1982). According to Radwan and El-Wakil (1991), cypermethrin-treated lettuce discs had the highest fatality percentages of any synthetic pyrethroid against *Eobania vermiculata* snails. Also, Amaeze *et al*. (2011) demonstrated that Cypercot with the active component cypermethrin had an immediate fatal impact on *Archachatina marginata* snails. On the other hand, lambda-cyhalothrin was the most effective, causing 70.0% of deaths in land snails, *M. cantiana*, Genena, and Mostafa (2008). Further, Lambada was shown to be less effective than methomyl on land snails, *Eobania vermiculata*, and *Helicella vestalis* by feeding utilizing a leaf dipping approach for seven days under lab circumstances by Hafez *et al*. (2022). Furthermore, it was the most potent agent against *Monacha cartusiana*, causing 100% death after 3 days compared to other insecticides (Hend *et al*., 2023). Moreover, it showed 100% death at 4, 6, and 8% concentrations in individuals after 3 days post-treatment (Maaroof *et al*., 2023).

*In vitro, Monacha obstructa* was found to be more sensitive to methomyl (lannate 90%) and had a very harmful impact by contact technique after 72 h of treatment (Hussein *et al*.,1999). As well, Aioub *et al*. (2000) demonstrated that methomyl was very poisonous and caused 100.0% of death in land snails, *M. cantiana*, Genena, and Mostafa (2008). Additionally, methomyl decomposed slowly in comparison to other pesticides tested against *M. cartusiana in vitro*, Ismail and Mohamed (2009). Otherwise, under field conditions, they found that methomyl had the greatest residual effect on *M. cartusiana* snails. So, it reduced *Monacha contiana* number of individuals more than using Vertimec (abamectin) in pea plantations, lettuce, and cabbage fields. Also, they observed that after one day of treatment with methomyl 2% wheat bran bait, the land snails examined were more effective. (Radwan and EL-Wakil, 1991; Abdallah *et al*., 1992; Ismail *et al*., 2005; Mortada *et al*., 2012 and Samy *et al*., 2015). Additionally Ali *et al*. (2020) From this study, it could be concluded that methomyl and other pesticides were efficient with the same dose to control tomato pests in Egypt as the chocolate banded snail, *E. vermiculata*. Moreover, Bashandy and Raddy (2021) indicated that methomyl was the most toxic compound against *M. obstructa* and the snails were more susceptible to methomyl because it caused 96% mortality for animals with LC_50_ values was 1.80% with leaf dipping techniques after seven days.

**In conclusion**, methomyl produced a much higher mortality in land snails than lambda. To test animals using touch methodology, it was more dangerous than lambda as a molluscicide than the leaf dipping method. Because of the hazards of pesticides on macroinvertebrates and human life, these pesticides must be used with caution, and their discharge into the terrestrial environment must be properly monitored and regulated.

## Acknowledgments

The authors would like to thank the Agricultural Zoology and Nematology Department, Faculty of Agriculture, Al-Azhar University, Cairo, Egypt, for providing the necessary laboratory for the experiment. Dr. Hany Bashandy (Department of Genetics, Cairo University, Giza, Egypt) provided useful comments on the work.

## Author contribution

A.S. Bashandy conceived and designed the experiment and contributed to writing the final manuscript. M.H. Awwad helped in the preparation of the experiment.

## Data availability

The article contains data.

## Additional information

There is no extra information for this publication.

## References

Abd El-All SM. 2004. Toxicity and biochemical response of Eobania vermiculata land snail to niclosamide molluscicide under laboratory and field conditions. J. Agric. Sci. Mansoura Univ., 29 (8): 4751–4756.

Abdallah EAM, Kassem FA and Kadous EA. 1992. Laboratory and field evaluation of local bait formulations of certain pesticides against mollusca species. J. Pest Control Environ. Sci., 46, 179–192.

Aioub AA, Ismail SAA and Mohamdein AA. 2000. Toxicological and histological studies on some pesticides-treated land snails. Proceeding of the first International Con. on Biological Science Faculty of Science Tanta Univ., 1 (2): 19:38.

Ali AM, El Roby ASM and Hassan HM. 2020. Controlling the Chocolate Banded Snail, Eobania vermiculata by Using some Insecticides at Minia Governorate, Egypt, J. of Plant Protection and Pathology, Mansoura Univ., 11(11):543–547.

Amaeze NH, Ugwoeje D and Egonmwan RI. (2011) Histopathological and physiological effects of selected agrochemicals on non-target Archachatina marginata. Yctijenvscs 1: 21–26.

Bakr EM. 2005. A new software for measuring leaf area, and area damaged by Tetranychus urticae Koch. Journal of Applied Entomology, 129 (3), 173–175.

Barker GM. 2002. Moulluscs as crop pests. CABI Publishing, CAB International, Walling ford, U.K., pp.: 468.

Bashandy AS. 2018. Studies On Some Terrestrial Gastropods Injurious to Some Important Crops at Giza Governorate. Ph.D. Thesis, Fac. Agric., Al-Azhar Univ., Egypt, 194 pp.

Bashandy AS and Raddy HM. 2021. In Vitro Assessment of The Efficacy of Metaldehyde, Methomyl and Copper Sulfate on some Terrestrial Gastropods. Journal of Plant Protection and Pathology, 12(7): 447–451.

El-Khodary AS, Sharshir FA, Helal RM and Shahawy, Wafaa A. 2001. Evaluation of some control methods against the land snail, Monacha cantiana (Montagu) at Kafr El-Sheikh Governorate, Egypt. J. Agric. Res. Tanta Univ., 27 (2): 290–300.

El-Okda MMK. 1979. Laboratory Studies on the Molluscicidal Toxicity of Methomyl and aldicarm against some land snails. Agric. Res. Rev., 57 (1): 199–207.

El-Wakil HB and Radwan MA. 1991. Biochemical studies on the terrestrial snail, Eobania vermiculata (Muller) treated with some pesticides. J. Environ. Sci. and Health, 26 (596): 479–489.

Eshra EH. 2014. Toxicity of methomyl, copper hydroxide and urea fertilizer on some land snails. Annals of Agricultural Science, 59(2), 281–284.

Genena MA and Mostafa FA. 2008. Molluscicidal activity of six pesticides against the two land snails, Monacha cantiana and Eobania vermiculata (Gastropoda: Helicidae) under laboratory conditions. Journal of Plant Protection and Pathology, 33(7), 5307–5315.

Genena Marwa AM. 2003. Studies on the terrestrial gastropods at Dakahlia Governorate. M.Sc. Thesis, Fac. Agric., Mansoura Univ., 136 pp.

Hafez ZM, Sobieha AK, Asran AA and Emam HM. 2022. Toxicity of Lambada Cyhalothrin, and Methomyl on Terrestrial Snails, Eobania vermiculata, and Helicella Vestalis, Under laboratory Conditions. Journal of Environmental Science, 51(4), pp. 93–108.

Hammond RB, Smith SS and Beek T. 1996. Timing of molluscicide application for reliable control in no-tillage field crops. J. Econ. Entomol., 89: 1028–1032.

Hanafy AHA, Youssef HM and El-Shahat MS. 1998. Preparation of methomyl baits and efficacy against certain land Mollusca in different vegetation. Adv. Agric. Res., 3 (3): 435–441.

Heiba FN, Al-Sharkawy MI and Al-Betal AA. 2002. Effect of the insecticide, lannate, on the land snails, Eobania vermiculata and Monacha cantiana, under laboratory conditions. J. Biological Sci., 2 (1): 8–13.

Heiba FN, Mortada MM, Geassa SN, tlam AI and Sahar IA. 2018. Terrestrial gastropods: Survey and relationships between land snail assemblage and soil properties. J. Plant Prot. and Path., Mansoura Univ., 9(3): 219–224.

Hussein HI, Al-Rajhy D, El-Shahawi FI and Hashem SM. 1999. Molluscicidal activity of Pergularia tomentosa (L.), methomyl and methiocarb, against land snails. International Journal of Pest Management, 45(3), 211–213.

Iglesias J, Castillejo J and Castro R. 2003. The effects of repeated applications of the molluscicide metaldehyde and the biocontrol nematode Phasmarhabditis hermaphrodita on molluscs, earthworms, nematodes, acarids and collembolans: a two_□year study in north_□west Spain. Pest Management Science: Formerly Pesticide Science, 59(11), 1217–1224.

Ismail SA, Abd-Allah SA, El-Massry SA and Hegab AM. 2005. Evaluation of certain chemicals and insecticides against Monacha cartusiana snails infesting some vegetable crops at Sharkia, Governorate. J. Agric. Sci. Mansoura Univ., 30 (10): 6283–6292.

Ismail SA, Shetaia SZS and Khattab MM. 2015. Time of application as main factor affecting the efficacy of certain pesticides against land snail Monacha cartusiana (Muller) under filed conditions at Sharkia Governorate. Journal of Plant Protection and Pathology, 6(5), 853–858.

Ismail SAA and Mohamaed DMO. 2009. Persistence of fresh prepared baits of certain pesticides tested at different intervals against Monacha cartusiana snails under laboratory conditions. Egypt. J. Appl. Sci., 24 (1): 274–280.

Kassab A and Daoud H. 1964. Notes on the biology and control of land snails of economic importance in the UARJ Agric. Res. Rev., 42, pp.77–98.

Maaroof HM, El Massry SA and Hassan AEA. 2023. Molluscicidal activity of Four Pesticides against the Glassy Clover Snail, Monacha Cartusiana under Laboratory and Field Conditions, a Comparative Study. Bulletin of Faculty of Science, Zagazig University, (1), pp. 176–180.

Mobarak SA. 2003. Mollusicidal activity of Methomyl and Diazinon against Land snail: Monacha obstructa and Eobania vermiculata. MSc. Fac. of Agric., Cairo University. pp: 116.

Mortada MM, Mourad AAM, Abo-Hashem AM and Keshta TMS. 2012. Land snails attacking pea fields: 1-Efficiency of certain biocides and molluscicides against Monacha sp. land snails at Dakahlia Governorates. J. Plant Prot., & Path., Mansoura Univ., 3: 717 –723.

Mourad AA. 2014. Molluscicidal effect of some plant extracts against two land snail species, Monacha obstructa and Eobania vermiculata. Egypt Acad. J. Biol. Sci., 6(1):11–6.

Ohayo GJA, Heederik DJJ, Kromhout H, Omondi BEO and Boleij JSM. 1997. Acetylcholinesterase inhibition as an indicator of organophosphate and carbamate poisoning in Kenyan agricultural workers. International Journal of Occupational and Environmental Health, 3: 210–220.

Radwan MA, and El-Wakil HB. 1991. Impact of certain carbamate and synthetic pyrethroid insecticides on the non-target terrestrial snail Eobania vermiculata. Alex. Sci. Exch., 12 (2): 305–316.

Samy MA, Fakharany SK M and Hendawy AS. 2015. Population fluctuation and host preference of land snail, Monacha spp and its control of biocides compared with neomyl. Fifth Intern. Comp. Plant Prot. Res. Inst. Hurgada. Egypt 3-9 May 2015. sustainable Agricultural Development the Agricultural Production and the Challenges of Plant Protections.

Vijverberg HP and van den Bercken J. 1982. Action of pyrethroid insecticides on the vertebrate nervous system. Neuropathol. Appl. Neurobiol., 8 (6):421–440.

Wafaa AS, Nadia MM and Hend S. El-Tahawe. 2018. Population Density, Food Consumption and Damage Caused by the Land Snail Monacha cantiana to Some Vegetable Crops at Kafr El-Sheikh Governorate. J. Plant Prot. and Path., Mansoura Univ., 9 (9): 601–604.

Zedan HA, Mortada MM and Shoeib, Amera A. 2006. Assessment of molluscicidal activity of certain pesticides against two land snails under laboratory and field circumstances at Dakahlia Governorate. J. Agric. Sci. Mansoura Univ., 31 (6): 3957–3962.

